# Validating a high-throughput *in vitro* model for culturing antibiotic-altered microbiome communities

**DOI:** 10.64898/2026.09.16.752229

**Authors:** Weimiao Long, Avani Tantry, Julielam Tran, Erika L. Cyphert

## Abstract

The gut microbiome plays a central role in host health, and its disruption is associated with metabolic, infectious, and immune conditions. Stool sampling is frequently used to characterize snapshots of the gut microbiota. To screen microbiota communities *in vitro*, multi-stage bioreactors have been developed that model distinct physiological environments along the gastrointestinal tract. However, such bioreactors are often operationally complex and provide limited throughput. Stool-derived *in vitro* communities offer several advantages to study microbiota due to their ease of use, high-throughput nature, and have previously been shown to remain stable following repeat passaging. Yet, prior characterization of stool-derived cultures has relied on 16S rRNA sequencing of unaltered gut microbiota, leaving several key questions unanswered: 1) how closely does *in vitro* culture recapitulate the functional potential and resistome composition of its inoculum and 2) can these cultures model antibiotic disrupted microbial communities? In this work, we generated stool-derived *in vitro* cultures from male and female mice with both unaltered and antibiotic altered gut microbiomes and applied shotgun metagenomic sequencing to characterize taxonomic composition, functional pathway capacity, and antimicrobial resistance gene (ARG) content. Antibiotic-altered microbiota communities, dominated by *Enterobacteriaceae*, were faithfully recapitulated in culture with preservation of both taxonomy and ARGs. By contrast, unaltered communities underwent substantial restructuring in culture, with *Bifidobacterium* and *Enterococcus* blooming and driving sex-divergent shifts in functional capacity and a substantial amplification of the resistome. Together, these findings validate a high-throughput stool-derived *in vitro* culture as a tractable proxy for an antibiotic altered gut microbiome to screen interventions.

**Importance:** When the gut microbiota is disrupted, patients face an elevated risk of infection, metabolic disease, and immune dysfunction. Restoring a disrupted microbiome community is challenging, in part due to a lack of models that represent disrupted communities to allow for rapid testing of interventions. Prior studies developed an *in vitro* stool-derived culturing platform for microbiota communities but did not evaluate the functional potential or antimicrobial resistance gene burden of these communities. In this work, we extend the characterization of this platform using shotgun metagenomic sequencing and show that it faithfully recapitulates the composition, function, and antimicrobial resistance gene content of antibiotic-disrupted microbiota communities. This platform provides a practical tool for evaluating disrupted gut microbiota and for screening interventional compounds.

## Introduction

The gut microbiota is a complex and dynamic community of microorganisms that is closely linked to the host’s health^1^. Yet, characterizing the gut microbiota poses inherent challenges. Since microbes have distinct biogeography in the gastrointestinal tract, sampling strategy strongly influences the microbial composition observed^2^. Prior studies have demonstrated that the sampling method significantly shapes the resulting compositional profile^3^. Despite spatial heterogeneity of gut microbiota along the gastrointestinal tract, stool sampling remains the most common approach in gut microbiome studies, owing to its accessibility and non-invasiveness^4^.

*In vitro* gut microbiome models have emerged as a tool to longitudinally study the dynamics of gut microbes and span from highly engineered multi-stage bioreactors^5–8^, to batch cultures^9–12^, to microfluidic gut-on-a-chip systems^13–15^. Although multi-compartmental bioreactor systems are powerful, they generally require specialized equipment and precise computer-controlled systems, limiting their accessibility and throughput. Batch culture offers a more high-throughput approach that can be readily implemented and regulated in a lab setting. Stool-derived *in vitro* communities (SICs) are an example of a batch culture platform and are generated by inoculating fecal samples into anaerobic cultures and serially passaged in rich media^10^. SICs have been shown to preserve much of the taxonomic diversity of the original stool inoculum in both mice and humans and remain stable and reproducible across replicates and passaging^10,16,17^.

Alteration of the gut microbiome can arise from many sources, including diet, disease, and antibiotic use^18^. Among these, antibiotic exposure is one of the most clinically relevant and well-characterized drivers of dysbiosis. Antibiotic use poses the risk of contributing to the development of *Clostridioides difficile* infection and antibiotic-induced microbiota changes are associated with obesity and diabetes among other conditions^19,20^. Beyond depleting diversity, antibiotic use also imposes antibiotic resistance. The collection of all antimicrobial resistance genes (ARGs) in a microbial community is termed the “resistome” and antibiotic use can increase the burden of the resistome^21^. The U.S. Centers for Disease Control and Prevention (CDC) estimates approximately 2.8 million resistance-associated infections annually in the United States alone and antibiotic resistance has become an urgent public health threat^22^. Therefore, understanding antibiotic-altered gut microbiota is of substantial interest.

While SICs were previously characterized, it is important to note that there are several remaining questions. First, prior characterization relied on 16S rRNA sequencing, that resolves taxonomic identity at the genera-level but provides limited information on the functional potential capacity of the community. Second, prior *in vitro* cultures focused on using stool from unaltered gut microbiota rather than antibiotic-disrupted systems. Here we sought to validate the *in vitro* SIC model generated from both unaltered and antibiotic-altered gut microbiome communities as a reproducible model to study the microbial dynamics at taxonomic, functional potential, and antibiotic resistance gene level. Specifically, we generated SICs from mice with both an antibiotic altered and unaltered microbiome community and used shotgun metagenomic to assess how well the model captured the taxonomic composition, functional potential, and antibiotic resistance gene composition relative to the starting stool samples. Our *in vitro* culture system recapitulated the antibiotic-altered gut microbiota closely. In accordance with prior work^10^, we found that recapitulation broke down when the starting inoculum community had higher diversity. Collectively, our findings position this culture platform as a tractable *in vitro* proxy for an antibiotic-altered gut microbiome for downstream screening of interventions.

## Results

### In vitro culture closely recapitulates an antibiotic-altered gut microbiome

Mouse fecal samples were first inoculated into brain heart infusion (BHI) media and later passaged using a 96-well plate format (**Figure 1A**). To account for potential sex-specific differences in response to antibiotic-driven alteration, we stratified both the Altered and Unaltered treatment groups by sex (female and male). We first sought to analyze how closely the *in vitro* culture model recapitulated the Altered gut microbiota relative to the initial stool sample. To visualize community composition at the genera taxonomic level, we identified the top 15 genera with mean relative abundance greater than 1% across all samples. These are depicted individually in stacked bar charts, with remaining genera at lower abundances summed into an “Others” category (**Figure 1B**).

**Figure 1.**
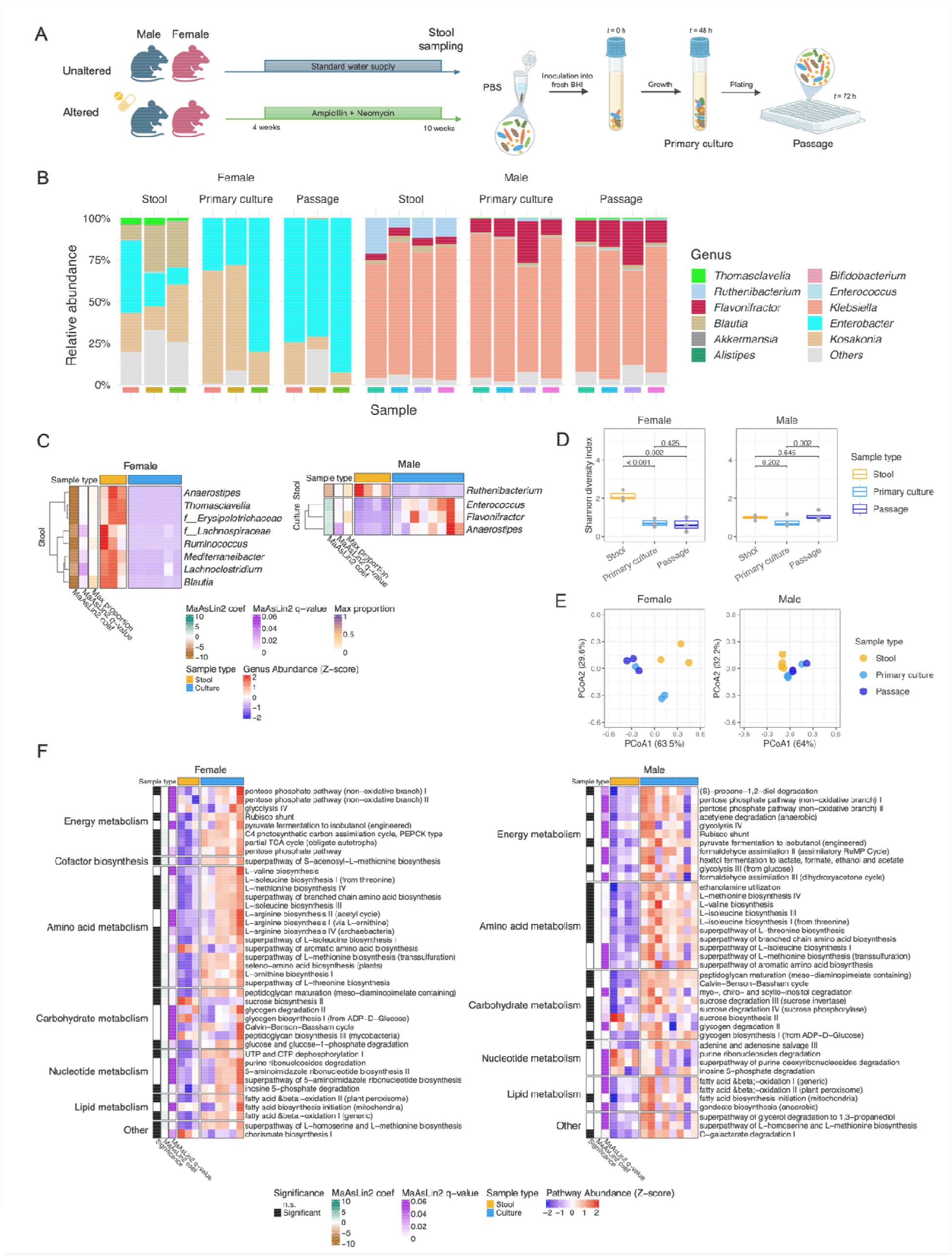
(A) Experimental setup. Fecal samples from mice with both Altered and Unaltered microbiota were inoculated into anaerobic batch culture and passaged. (B) Taxonomic composition of microbial communities at genera level in the Altered group. Stool samples, *in vitro* primary culture, and passage are depicted. The color under each stacked bar represents an individual mouse replicate. (C) Heatmap of differentially abundant genera between stool and culture in the Altered group (female and male). Feature values were normalized using row-wise Z score transformation (mean = 0, standard deviation = 1) to emphasize relative differences across samples. Rows represent the genera that were significantly enriched in either group (MaAsLin2 q-value < 0.05; max proportion > 1%). (D) Boxplot comparing Shannon diversity index of genera level composition between stool, primary culture, and passage. (E) Principal coordinate analysis (PCoA) based on the Bray-Curtis distance metric calculated on genera level composition between stool, primary culture, and passage. (F) Heatmap of 40 most abundant functional potential pathways between stool and culture in females (left) and males (right) from the Altered group.

The Altered microbiota demonstrated sex-dependent structure both in stool and in culture. Female Altered stool samples contained a mixed community dominated by opportunistic pathogens of *Kosakonia* and *Enterobacter* (approximately 25% average relative abundance for each), with persistent commensal *Firmicutes* including *Blautia* and *Thomasclavelia* present at moderate abundance. Male Altered stool samples, by contrast, were dominated by the opportunistic pathogen *Klebsiella* (nearly 75% relative abundance across all stool samples), with *Flavonifractor*, *Blautia*, and *Ruthenibacterium* present as minor constituents. The *in vitro* culture derived from Female Altered stool samples recapitulated the dominant *Enterobacterales* taxa (*Kosakonia* and *Enterobacter*) but lost some residual commensal *Firmicutes*. Cultures derived from Male Altered stool samples nearly completely preserved *Klebsiella* and other key taxa found in the stool. To investigate the effect of passaging once the microbial community was in culture, we performed differential abundance testing using MaAsLin2 (Microbiome Multivariable Associations with Linear Models)^23^. Specifically, we sought to identify taxa that significantly differed between each sex-specific primary culture and subsequent passage. Within the Altered groups, we found that *Enterobacterales* (*Kosakonia*, *Enterobacter*, *Klebsiella*) were preserved at similar relative abundances in the passage relative to their respective primary cultures (**Supplemental Figure 1**). While we did detect some subtle differences between the taxonomy of the primary culture and passage in the Male Altered group, overall the community structure was largely maintained.

We also used MaAsLin2 to identify taxa that significantly differed between stool and culture for each sex. To increase statistical power, the primary culture and passage were pooled together for each sex into a single “culture” group for this statistical analysis. The sample pooling was justified by the compositional consistency between the primary culture and the passage. In the Female Altered group, 29 genera were significantly enriched in stool (MaAsLin2 q-value < 0.05), and none were enriched in the culture. In the Male Altered group, 68 genera were enriched in stool and 5 in the culture. Notably a large percentage of these significantly differentially abundant genera were at a relative abundance < 1% (72% in females, 95% in males). To identify the taxa that were both differentially abundant and had a relative abundance > 1%, we filtered the differentially abundant taxa by their maximum relative abundance and visualized the results in heatmaps (**Figure 1C**). Across both sexes, the *in vitro* cultures derived from the Altered microbiomes closely recapitulated the original taxa found in stool with only a few taxa shifting. In the Female group, we found the main constituents of the stool (*Kosakonia* and *Enterobacter*) were retained at a similar abundance in culture, though commensal *Firmicutes* were depleted in culture. In the Male group, the main genus (*Klebsiella*) was retained at a similar abundance from stool to culture with other microbes being depleted or enriched (*Ruthenibacterium* depleted in culture and *Enterococcus*, *Flavonifractor*, and *Anaerostipes* enriched in culture). Across both sexes, the genera that experienced the greatest changes from stool to culture all belonged to the phylum *Firmicutes* (83% in females, 77% in males).

To assess whether overall community structure was retained from stool to culture, we calculated the alpha and beta diversity. Shannon diversity (calculated at the genera level) indicated that after 48 hours of culture, the diversity significantly decreased in females (p < 0.001), but not in males (p = 0.20) (**Figure 1D**). This indicated that while the composition of the most dominant taxa was largely retained from stool to culture, that there was a loss of some taxa in culture, particularly in the Female Altered group. Moving from primary culture to passage, Shannon diversity was comparable in the Female Altered group (p = 0.425) but significantly decreased in the Male Altered group (p = 0.002). We identified 19 significantly differentially abundant genera between the primary culture and passage using MaAsLin2 in the Male Altered group, though the majority of these genera (84%) were at relative abundance < 1% (**Supplemental Figure 1**). *Klebsiella* was the only genera that was depleted in the passage relative to the primary culture and it changed in average relative abundance from 80% to 71%.

Bray-Curtis beta diversity was performed on the genera level to identify differences in community structure for each sex between the stool, primary culture, and passage (**Figure 1E**). Overall, the Male Altered group demonstrated tighter clustering of cultured samples to stool compared to the Female Altered group. PERMANOVA (Permutational Multivariate Analysis of Variance) tests were performed within each sex with permutations restricted within mouse to account for the paired design. Resulting p-values were interpreted with caution given the small sample size and permutation ceiling.

To investigate how the compositional shift from stool to culture influenced the functional potential capacity of the community, we performed MaAsLin2 on the functional pathways profiled (**Supplemental Table 1**). In the Female Altered group, we identified 222 differentially abundant pathways between stool and culture and there were 127 differentially abundant pathways in the Male Altered group. We then visualized the 40 most abundant pathways (measured by average abundance across all samples compared, excluding UNMAPPED and UNINTEGRATED), annotated by MaAsLin2 significance (q-value threshold at 0.05) and direction of enrichment (**Figure 1F**). Amino acid and carbohydrate metabolism were the most prominent ontology classes that were significantly enriched in culture relative to stool.

### In vitro culture restructures the Unaltered gut microbiome at taxonomic and functional level

We next examined how the *in vitro* culture model recapitulated the Unaltered gut microbiome from stool to culture. At the genera level, stool samples from the Female Unaltered and Male Unaltered groups shared a similar, diverse community containing 50-75% of *Bacteroidetes* (*Duncaniella*, *Alistipes*, and *Muribaculum*) (**Figure 2A**). The “Others” category included all low abundance but detectable taxa and accounted for 20-50% of reads in stool, reflective of the high diversity of the unperturbed mouse gut. When placed in culture, this diversity collapsed almost completely with depletion of *Bacteroidetes* to low or undetectable abundance that diverged by sex. Cultures derived from Female Unaltered mice were dominated by *Enterococcus* in two out of four replicates and by *Bifidobacterium* in the remaining replicates, with all cultures retaining *Akkermansia*. Cultures derived from Male Unaltered mice were also heterogeneous. Specifically, *Enterococcus* and *Bifidobacterium* dominated and *Klebsiella* colonized in two replicates. Low but detectable levels of *Akkermansia, Thomasclavelia*, and *Clostridioides* were minor constituents. Microbial communities observed in primary cultures were preserved at nearly identical relative abundances in the passages, indicating that the community remained stable once it reached culture (**Figure 2A**, **Supplemental Figure 2**). Taxa that significantly differed between stool and *in vitro* culture were determined separately by sex using MaAsLin2. At the genera level, volcano plots showed widespread taxonomic collapse in the culture relative to the stool in both sexes (**Figure 2B**). Only three genera thrived significantly (q < 0.05) *in vitro* that split by sex. *Bifidobacterium* bloomed in both sexes, while *Akkermansia* and *Enterococcus* expanded in females and males, respectively.

**Figure 2.**
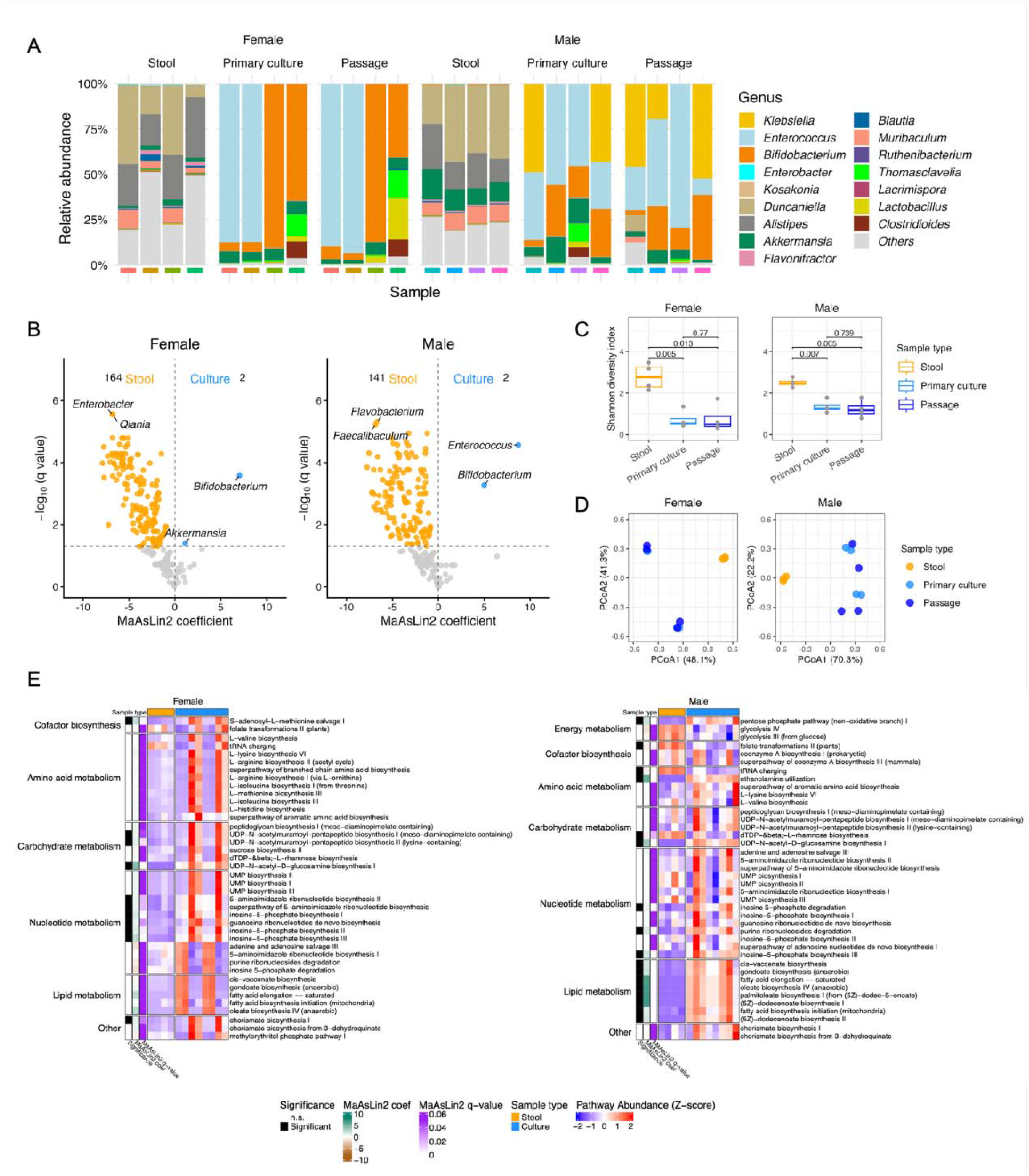
(A) Taxonomic composition of microbial communities from stool, primary culture, and passage of the Unaltered group at the genera level. (B) Volcano plots of differentially abundant genera between stool and culture in the Female (left) and Male (right) Unaltered groups. Genera significantly enriched in stool (q < 0.05) are in orange, those significantly enriched in culture are in blue, and non-significant genera are in grey. (C) Boxplot comparing Shannon diversity index of genera composition between stool, primary culture, and passage. (D) Bray-Curtis beta diversity analysis of genera composition between stool, primary culture, and passage. (E) Heatmap of the 40 most abundant metabolic pathways characterized by HUMAnN3 between stool and culture in Female (left) and Male (right) Unaltered group.

To assess how community structure shifted from stool to culture in Unaltered samples, we examined both the alpha and beta diversity. Relative to Unaltered stool samples, the Shannon diversity significantly decreased in primary culture in both males and females (p < 0.01). However, the alpha diversity in both sexes remained stable once it reached culture (p > 0.05), suggesting that passaging had minimal impact on the diversity of the community. Overall, stool samples from the Unaltered group showed a significantly higher diversity than stool samples from the Altered group (Welch’s t-test, p = 0.002), consistent with the alteration that was applied *in vivo* (**Supplemental Figure 3**). The ordination distance between stool and culture samples on the Bray-Curtis PCoA plot for the Unaltered group was substantially greater than the Altered group (**Figure 2D**), suggesting that the compositional drift in culture was exacerbated when the starting inoculum was more diverse. Statistical significance of community compositional differences was assessed within each sex using PERMANOVA. Resulting p-values were interpreted with caution given the small sample size and permutation ceiling.

To investigate how the compositional shift from stool to culture influenced the functional potential capacity of the community, we performed differential abundance analysis on the functional pathways profiled by HUMAnN3^24^. The Female Unaltered group had 138 significantly differentially abundant functional pathways between stool and culture and the Male Unaltered group had 109. We then visualized the 40 most abundant pathways annotated by MaAsLin2 significance (q-value threshold at 0.05) and direction of enrichment (**Figure 2E**). The Unaltered group showed a cluster of culture-enriched pathways emerging in both sexes. In the Female Unaltered group, nucleotide biosynthesis pathways were the most significantly enriched category, consistent with the Bifidobacterial bloom in culture^25^. In the Male Unaltered group, lipid metabolism pathways were the dominant culture-enriched category including cis-vaccenate biosynthesis and anaerobic fatty acid biosynthesis, consistent with *Enterococcus* colonization in culture^26^. Together, these two categories represent the biosynthetic demands of fast-growing bacteria and their enrichment matches the metabolic needs of *Bifidobacterium* and *Enterococcus* blooms in culture (**Figure 2A**).

### In vitro culture recapitulates the resistome of an antibiotic altered gut microbiota

To determine how well our *in vitro* culturing platform of the Altered microbiome community recapitulated the antimicrobial resistance gene composition of the stool, we analyzed both the richness and total burden of resistance genes in the resistome. We calculated total ARG burden as the sum of normalized depth per million (dpm) across all detected ARGs per sample. Female Altered stool samples had an average ARG burden of 85.5 ± 10.8 dpm and 143.4 ± 9.8 dpm in Male Altered stool samples, which indicated that the Altered stool had a relatively high resistance gene burden prior to culturing. Across stool and culturing, the total ARG burden did not significantly differ in Altered microbiome samples, independent of sex (**Figure 3A**; female p = 0.14, male p = 0.24, paired t-test). ARG richness (number of distinct ARGs detected per sample) had an average of 48.0 ± 3.6 in Female Altered stool and 54.2 ± 1.9 in Male Altered stool, reflecting the large diversity of resistance genes already present in the Altered microbiome communities. As with total ARG burden, ARG richness was preserved across stool to culturing in Altered microbiome samples (**Figure 3B**; female p = 0.12, male p = 0.74, paired t-test). Further, both total ARG burden and richness were preserved from primary culture to passage in both sex groups. Preservation of both burden and richness of ARGs in Altered microbiome samples indicated that the *in vitro* culture was able to maintain the resistome state in Altered stool samples.

**Figure 3.**
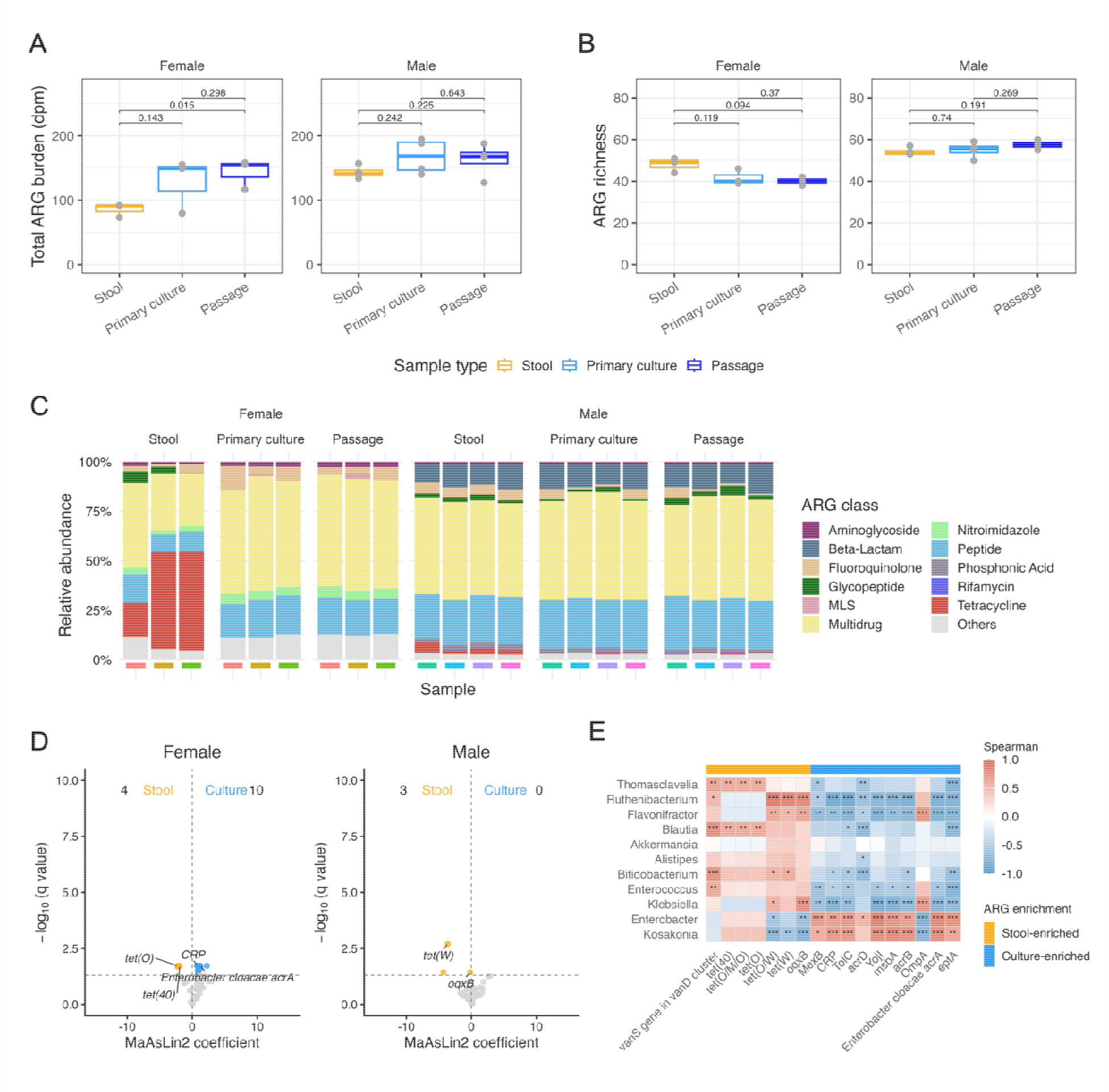
(A) Total ARG burden and (B) ARG richness amongst stool, primary culture, and passage in antibiotic Altered samples, faceted by sex. (C) Resistome composition of stool, primary culture, and passage of the antibiotic Altered group. (D) Volcano plots of differentially abundant ARGs between stool and culture in the Female (left) and Male (right) Altered groups. ARGs significantly enriched in stool (q < 0.05) are in orange, ARGs enriched in culture are blue, and non-significant ARGs are in grey. (E) Spearman correlation matrix between genera relative abundance (rows) and ARG relative abundance (columns) in Altered samples. Rows were restricted to genera with mean relative abundance > 1%. Columns are restricted to ARGs identified as differentially abundant between stool and culture by MaAsLin2 (q < 0.05). Asterisks indicate significance (*p < 0.05, **p < 0.01, ***p < 0.001, Benjamini-Hochberg correction FDR).

We then investigated if the resistome composition was preserved. Specifically, we calculated the per-sample relative abundance of each ARG drug class and visualized the results in stacked bar plots (**Figure 3C**). Across both sexes, the dominant ARG classes in Altered stool samples were multidrug, peptide, beta-lactam, and fluoroquinolone resistance, with female stool additionally harboring a moderate fraction of tetracycline class ARGs (approximately 30% relative abundance). The resistome composition in cultures from the Male Altered group was visually indistinguishable from the stool samples. Whereas the resistome composition in cultures from the Female Altered group largely conserved the composition of the stool sample but demonstrated a modest loss in tetracycline-associated resistance genes. Differential abundance analysis was performed using MaAsLin2 to find specific ARGs that significantly differed between stool and in culture (**Figure 3D**). Across both sexes, only a small number of ARGs reached significance, indicating that the resistome composition of the Altered stool was largely recapitulated *in vitro* across both sexes.

Upon characterizing the resistome composition of the Altered microbiome samples from stool to culture, we next sought to evaluate whether associations between taxa and ARGs could mechanistically explain the microbial dynamics that we observed from stool to culture. To evaluate this, we selected the most abundant genera (mean relative abundance ≥ 1% as in **Figure 1B** and **Figure 2A**) and ARGs that significantly differed from stool to culture (MaAsLin2 q < 0.05, pooling both sexes) for our correlation analysis. Spearman rank correlations were then computed across all Altered samples and visualized in a correlation matrix (**Figure 3E**). Across both sexes, the stool-enriched ARGs consisted almost entirely of tetracycline class ARGs (*tet(40), tet(O), tet(O/M/O), tet(O/W), tet(W)*) together with *vanS* (histidine protein kinase from vanD vancomycin resistance cluster), and *oqxB* (multidrug efflux gene)^27,28^. These ARGs showed significant positive correlations with a set of commensal *Firmicutes* (*Blautia, Ruthenibacterium*, and *Thomasclavelia*), consistent with previous findings^29,30^. Their co-occurrence may help to explain the loss of both *Firmicutes* and tetracycline-associated ARGs in culture.

### In vitro culture amplified and restructured the resistome of Unaltered gut microbiota

To determine how well *in vitro* cultures of the Unaltered microbiome community recapitulated the antimicrobial resistance gene composition of stool, we analyzed the richness and total burden of resistance genes in the resistome. Female Unaltered stool had an average total ARG burden of 7.3 ± 3.0 dpm and Male Unaltered stool had 8.3 ± 3.3 dpm, which was consistent with the relatively low resistance gene burden expected of an unperturbed mouse gut microbiome. After culturing, total ARG burden rose to an average of 128.9 ± 79.7 dpm in females and 150.3 ± 37.6 dpm in males (**Figure 4A**). The increase was statistically significant in males (p = 0.005, paired t-test) and marginally significant in females (p = 0.054, paired t-test). We hypothesized that the female result was likely partially attributable to the heterogeneity of the changes in taxa from stool to culture (**Figure 2A**). Notably, ARG richness did not significantly change from stool to culture in either sex, despite the dramatic increase in ARG burden (**Figure 4B**; female p = 0.20, male p = 0.12, paired t-test). Total ARG burden and richness generally remained stable from primary culture to passage in Unaltered cultures of both sexes (with the exception of Female ARG richness, p = 0.014), demonstrating that in general passaging did not substantially perturb resistome dynamics.

**Figure 4.**
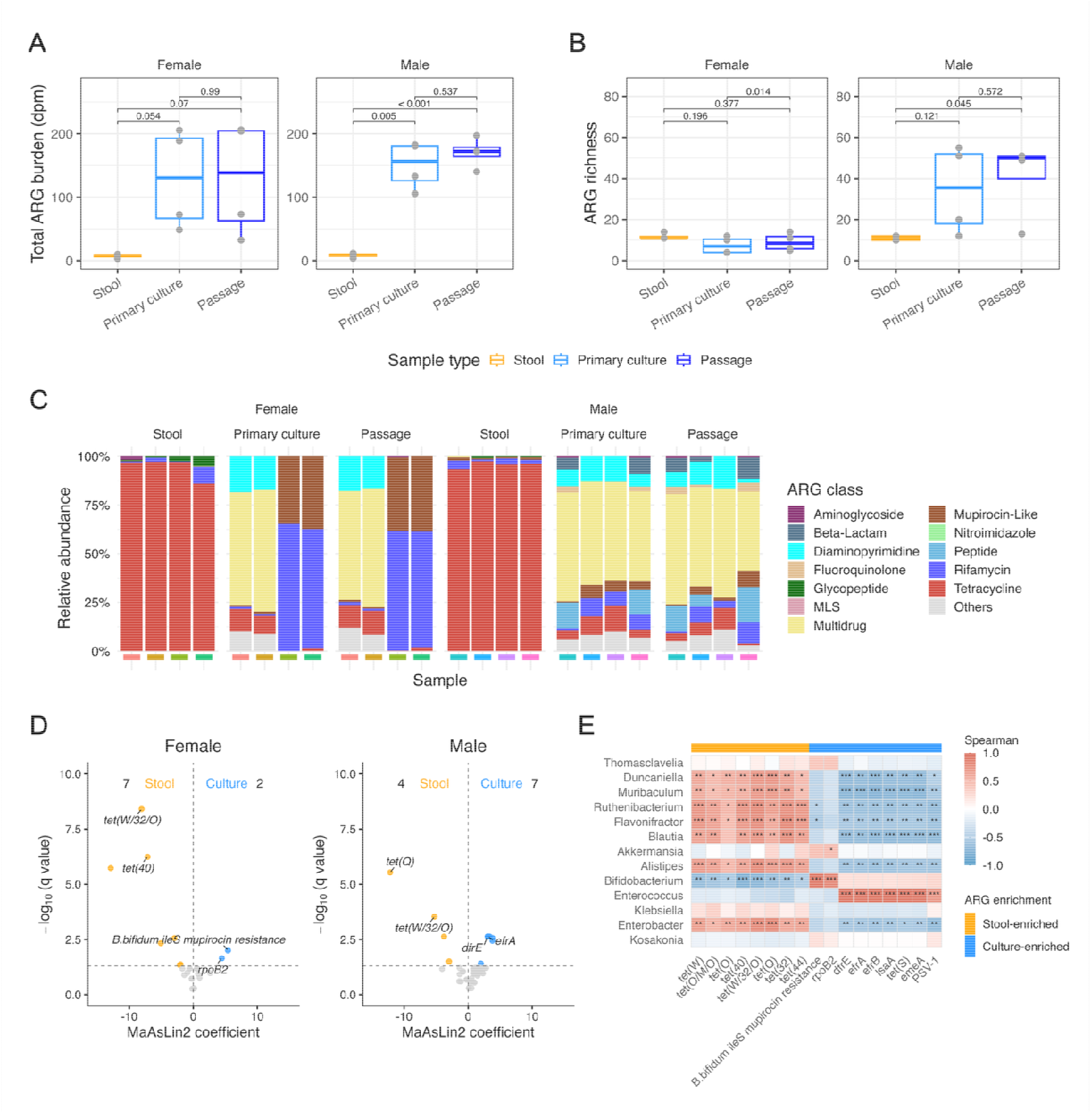
(A) Total ARG burden and (B) ARG richness in stool, primary culture, and passage in Unaltered samples, faceted by sex. (C) Resistome composition of microbial communities of stool, primary culture, and passage of Unaltered samples. (D) Volcano plots of differentially abundant ARGs between stool and culture in the Female (left) and Male (right) Unaltered groups. ARGs significantly enriched in stool (q < 0.05) are in orange, ARGs enriched in culture are in blue, and non-significant ARGs are in grey. (E) Spearman correlation matrix between genera relative abundance (rows) and ARG relative abundance (columns) in Unaltered samples. Rows were restricted to genera with a mean relative abundance > 1%. Columns were restricted to ARGs identified as differentially abundant between stool and culture by MaAsLin2 (q < 0.05). Asterisks indicate significance (*p < 0.05, **p < 0.01, ***p < 0.001, Benjamini-Hochberg correction FDR).

We also visualized per-sample relative abundance of ARG classes in stool and culture (**Figure 4C**). Unaltered stool samples were dominated by tetracycline-associated ARGs (approximately 95% across all stool samples in both sexes). This trend was consistent with the Unaltered stool being dominated by *Bacteroidetes* and *Firmicutes*, which are largely associated with tetracycline ARGs ^29,30^ (**Figure 2A**). After culturing, this dominance of tetracycline-associated ARGs largely collapsed and was replaced by a heterogeneous mix of ARG classes that varied across replicates and between sexes. The composition of the resistome was largely retained from primary culture to passage, demonstrating that the resistome composition generally remained stable once in culture.

As was observed in taxa, culture samples in the Female Unaltered group split into two compositional patterns of ARGs. Specifically, two replicates were enriched with multidrug and diaminopyrimidine resistance classes, while the other two were enriched in rifamycin and mupirocin-like resistance. In the Male Unaltered group, cultures were more consistent in composition, with multidrug (51.3% ± 5.5%), peptide (12.7% ± 4.1%), diaminopyrimidine (9.9% ± 4.6%), tetracycline (6.8% ± 4.2%), rifamycin (6.1% ± 3.8%), and beta-lactam (4.3% ± 4.3%) classes all present at moderate abundance. This compositional restructuring of ARGs mirrored the taxonomic transition from stool to culture described earlier. Differential abundance testing on individual ARGs was performed using MaAsLin2 and visualized as volcano plots (**Figure 4D**). In Female Unaltered samples, stool was enriched with tetracycline-associated ARGs (*tet(Q), tet(40), tet(W), tet(W/32/O)*), whereas culture was enriched with ARGs associated with rifamycin and mupirocin (*rpoB2*, *Bifidobacterium bifidum ileS conferring resistance to mupirocin*). Similarly, in Male Unaltered samples, stool was enriched with tetracycline-associated ARGs (*tet(Q), tet(44), tet(32), tet(W/32/O)*) and culture was enriched with more diverse ARGs (*efrB, efrA, dfrE, lsaA*).

To establish a mechanistic link between taxonomy and resistome composition for the Unaltered samples, we computed Spearman correlations on the genera and ARGs using the same strategy described earlier for the Altered groups (**Figure 4E**). *Bacteroidetes* (*Duncaniella, Muribaculum*) and *Firmicutes* (*Blautia, Flavonifractor*) were significantly positively correlated with tetracycline-associated ARGs. Additionally, *Enterococcus* was significantly positively correlated with *dfrE, efrA, efrB, lsaA, emeA, tet(S),* and *PSV-1*. Most of these ARGs are well-characterized intrinsic resistance genes in the genome of *Enterococcus*^31–34^. In the Male Unaltered group, both *Enterococcus* and *Bifidobacterium* were enriched in culture relative to stool. *Enterococcus* was significantly positively correlated with culture-enriched ARGs (*dfrE, efrA, efrB, lsaA, emeA, tet(S),* and *PSV-1*) specific to the Male Unaltered group, which pointed to *Enterococcus* primarily driving the restructuring of the resistome in culture. Additionally, *Bifidobacterium* was significantly positively correlated with two ARGs of particular interest. Specifically, *Bifidobacteria* have an intrinsically resistant form of ileS (isoleucyl-tRNA synthetase) that confers resistance to mupirocin that agreed with the correlation observed in our data^35^. Additionally, *rpoB2* is an RNA polymerase beta-subunit gene that confers intrinsic rifamycin resistance and is a well-characterized resistance gene in *Bifidobacteria*^36^. The strong positive correlation of these two genes with *Bifidobacterium* abundance was consistent with the observation that two Female Unaltered culture replicates with the highest *Bifidobacterium* relative abundance also showed the highest rifamycin and mupirocin-like class composition.

## Discussion

This work evaluated a high-throughput *in vitro* culture system, specifically stool-derived *in vitro* communities (SICs) as a model for studying gut microbial composition, functional potential, and resistome under conditions of an antibiotic-altered microbiota. We characterized the *in vitro* culture model at three different levels: taxonomic composition, functional potential capacity, and ARG composition. Across all three readouts, antibiotic Altered microbiome communities were closely recapitulated, while Unaltered microbiome communities underwent substantial restructuring driven by *Bifidobacterium* and *Enterococcus* bloomers.

We attribute the restructuring of the Unaltered community in culture to be partially linked to the media used and the difficulties of capturing the intricacies of the *in* vivo environment *in vitro*. Specifically, BHI broth is a liquid general-purpose culture media that contains infusions of brain and heart tissue and peptones to supply protein and other nutrients. Although BHI broth is shown to be capable of supporting the growth of fastidious and non-fastidious microorganisms, culture without host cells, mucin, or indigestible fiber inherently cannot fully recapitulate the gut environment. Therefore, some divergence in culture relative to the stool is expected and the community often collapses further as cross-feeding interactions are disrupted and ecological niches are depleted in culture^37^. This is particularly relevant for the Unaltered group in culture. The highly diverse Unaltered community requires host-derived substrates (e.g., mucin) and a structured physiological environment that culture cannot provide. A 3D *in vitro* model using mucin-coated scaffolds and culture demonstrated that mucin can shift the fecal microbiota towards an enrichment of mucus-associated bacteria *in vitro*^14^. Therefore, a greater divergence from the stool composition is expected in the Unaltered group relative to the Altered group, as we observed.

We hypothesize that specific taxa were enriched or depleted in culture relative to the stool samples for several reasons. Specifically, *Duncaniella* and *Muribaculum* are genera from *Muribaculaceae* whose metabolism relies on endogenous mucin glycans and exogenous polysaccharides like dietary fibers^38^. Since neither of these substrates were present in the *in vitro* culture, this likely explains the depletion of *Muribaculaceae*. Our findings were consistent with others that reported the depletion of *Muribaculum* in stool-derived Unaltered microbiome cultures^10^. Further, the genome of *Bifidobacterium* is particularly rich in carbohydrate-utilizing enzymes, making it highly capable of adapting to different carbon sources and colonizing and establishing a niche^39^. Notably, *Bifidobacterium pseudolongum*, a keystone species associated with Bifidobacterial blooms, was significantly enriched in both male and female Unaltered cultures relative to stool (**Supplemental Figures 4-5**)^40^. Moreover, the enrichment of *Enterococcus* in culture aligned with previous findings^12^. Interestingly, *Akkermansia* bloomed in cultures in Female groups and was maintained at a comparable relative abundance (slightly decreased, q-value > 0.05) in Male groups. This result was unexpected, given that *Akkermansia* is known to be heavily reliant on mucin as a carbon and nitrogen source^41^.

We further hypothesize that the heterogeneity of the microbial composition from stool to culture across replicates can be partly attributed to the “early bird” resource use dynamics described by Aranda-Díaz et al., in which the species that first consumes an available nutrient in a complex medium monopolizes the niche, even if that species was rare in the original inoculum^42^. Combined with the effects of dilution from stool to the *in vitro* culture, these resource use dynamics likely amplify small stochastic differences in the starting microbial composition, producing the divergent communities observed in fully grown cultures^10^.

The dissociation between the burden and richness of ARGs from stool to culture in the Unaltered microbiome samples potentially implied that *in vitro* culturing was amplifying the signal of the resistance genes that were already present at low abundance in stool samples. We hypothesize that this amplification of ARGs is mediated by microbes that are enriched in culture (e.g., *Enterococcus* and *Bifidobacterium*). Specifically, *Enterococcus* and *Bifidobacterium* expanded from < 1% relative abundance in stool to > 50% in culture and the ARGs that they carried (*rpoB2, dfrE, efrA,* and *efrB*) were correspondingly enriched in culture. *In vitro* culture amplified and restructured the resistome of Unaltered gut microbiota and taxonomic transitions appeared to drive the shifts in the resistome. Despite the microbial compositional shifts observed between stool and culture in the Unaltered group, the strong correlation between taxonomic and resistome composition opens future opportunities for studying specific gene families in defined synthetic communities cultured *in vitro*.

It is important to note several limitations of our work. Specifically, we acknowledge that this work is a preliminary investigative study on the reproducibility and screen capability of this *in vitro* culture system, and our small sample size inherently limited the statistical power for analyses. Future follow-up studies should include a more robust sample size to further validate our findings. Further, databases used for classification of ARGs are biased to well-characterized microbes. Additionally, pairing the existing shotgun metagenomic readout with metabolomics, metatranscriptomics, and PMA-treated sequencing would help to capture active responses, downstream metabolic outputs, and live versus dead cell distinctions that cannot be determined from existing datasets. Finally, using culturomics techniques to engineer the culture environment could help to recover microbial communities more faithfully. For example, by supplementing mucin, indigestible fiber, and bile acid mixes, the *in vitro* cultures could potentially rescue the mucus-associated and fiber-specialized commensals that were lost during culture. Extending this platform from mouse to human inocula, through humanized mouse models or patient fecal samples, would test whether the patterns observed translate across species.

## Materials and Methods

### Mouse strains

All animal use protocols were compliant with the Institutional Animal Care and Use Committee (IACUC) Office at UC San Diego. Male and female C57Bl/6J mice were purchased from Jackson Laboratory (Sacramento, CA, USA) and bred using trio breeding. Pups obtained from this breeding cycle were used for experiments. Litters were weaned at three weeks of age and randomly separated into cages by sex and treatment (n=5/cage). All cages contained corn cob bedding (Envigo Sani-Chip 7090A standard, CA, USA), standard laboratory chow (8604 Rodent Diet, Harlan Teklad, USA), deionized (DI) water *ad libitum* which was filtered and treated with bleach for sterilization, and cotton-based nesting material (Envigo Nestlets, CA, USA).

### Study design

Cohorts of male and female pups were each divided into Unaltered and Altered groups: Female Unaltered, Male Unaltered, Female Altered, Male Altered (n=5/group/sex, n=20 total). One Female Altered mouse was lost at eight weeks of age due to natural causes. Mice in the Altered group were dosed with an antibiotic cocktail consisting of 0.5 g/L neomycin (Sigma Aldrich, USA) and 1 g/L ampicillin (Sigma Aldrich, USA) administered via their drinking water, whereas the Unaltered group received standard drinking water. To prevent light-induced degradation of antibiotics, water bottles of the Altered group were wrapped in aluminum foil and further covered with polypropylene material. Mice in the Altered group received antibiotics starting at four weeks of age and received continuous perturbation to their microbiota until ten weeks of age. The antibiotic cocktail was replaced every 2-3 days over the duration of the study. Ampicillin and neomycin were selected due to their poor oral bioavailability, capable of acting directly on gut microbes while minimizing systemic effects as much as possible^29,43^.

### Fecal sampling

Fecal samples were collected from four randomly selected mice per group (except for Female Altered group that only had four mice) at the conclusion of the six week microbiota disruption period. Stool samples were collected and used for inoculating *in vitro* cultures and for direct shotgun metagenomic sequencing of full stool. Samples were collected from the same cohort of individual mice, yielding paired samples per mouse. For collection, individual mice were placed in empty sterile plastic containers and allowed to defecate normally. All samples were collected between 8:00 AM and 11:00 AM to minimize circadian effects on microbiota composition^44^.

#### In vitro culture inoculation

Fresh fecal samples were placed into 2.0 mL microcentrifuge tubes (Simport Scientific Cat. No. T339-6SPR), weighed, and transported into the anaerobic chamber within 30 minutes of collection.

#### Full stool

Fresh fecal samples were collected into sterile 2.0 mL cryovials (CellPro VC118-C Cryovials, Alkali Scientific, USA), weighed, and transported directly to the -80°C freezer within 30 minutes of collection. Samples were kept at -80°C until downstream shotgun metagenomic sequencing was performed.

### Stool-derived in vitro communities (SICs)

Stool-derived *in vitro* communities (SICs) were generated following the approach detailed in Aranda-Díaz et al.^10^, with modifications described below. To each 2.0 mL microcentrifuge tube containing fecal samples, 1 mL of anoxic PBS (Sigma Aldrich P4417-100TAB) with 0.05% (w/v) L-cysteine (Sigma Aldrich C5360-100G) was added, irrespective of fecal mass. Samples were vortexed (VWR Mini Vortex Mixer Cat. No. 10153-688) for three minutes to disintegrate large pieces of fecal matter and centrifuged (VWR Mini Centrifuge) for five minutes to separate the bacterial suspension from fibrous debris.

For the primary culture, 200 μL of the resulting supernatant was inoculated into 4 mL of anoxic brain heart infusion (BHI) medium (BD BBL) with 0.05% (w/v) L-cysteine in a 5 mL round-bottom tube (Corning FALCON). Tubes were capped and vented to allow gas exchange during incubation and placed on a tube rack and incubated in an anerobic chamber (Coy) at 37°C for 48 hours with continuous shaking (Boekel Scientific Variable Speed Mini Orbitron, speed dial 12). One primary culture was established per mouse (n=16).

Every primary culture was passaged in fresh anoxic BHI. Specifically, 150 μL primary culture was combined with 50 μL fresh medium per well in a 96 well plate (Avantor VWR Cat. No. 734- 2781), giving a final volume of 200 μL per well. This minimal dilution was intended to maintain a high bacterial density approximating the bacterial loads in the colon and to preserve accumulated metabolites^45^. Each primary culture was passaged in quadruplicate for downstream DNA extraction and in triplicate for optical density (OD) measurements. Additionally, 200 μL of fresh anoxic BHI medium without stool inoculum was added in triplicate to wells to serve as negative control and OD baselines. Plates were sealed with adhesive storage film (Eppendorf) perforated above each well with a sterile 20-gauge needle to permit gas exchange and incubated at 37°C for 24 hours with continuous shaking (speed dial 4).

Samples were collected for shotgun metagenomic sequencing at two stages. The remaining volume of each primary culture was transferred to a 2.0 mL cryovial (Alkali Scientific CellPro) and stored at -80°C. Following 24 hours of culturing in the 96-well plate, four replicate wells of each SIC were pooled, per condition, into a single 2.0 mL cryovial to obtain sufficient DNA, sealed, and stored at -80°C until DNA extraction. Optical density at 600 nm was measured on a plate reader (SpectraMax iD3, Molecular Devices) immediately after incubation. Background optical density from blank wells (fresh anoxic BHI without stool inoculum) was subtracted from all measurements.

### Shotgun metagenomic sequencing

The UC San Diego Microbiome Core performed nucleic acid extractions using previously published protocols^46^. Briefly, samples were purified using the MagMAX Microbiome Ultra Nucleic Acid Isolation Kit (Thermo Fisher Scientific, USA) and automated on KingFisher Flex robots (Thermo Fisher Scientific, USA). Blank controls and mock communities (Zymo Research Corporation, USA) were included per extraction plate, which were carried through all downstream processing steps. DNA was quantified using a PicoGreen fluorescence assay (Thermo Fisher Scientific, USA) and metagenomic libraries were prepared with KAPA HyperPlus kits (Roche Diagnostics, USA) in a miniaturized 1/5^th^ reaction volume format, and automated on EpMotion automated liquid handlers (Eppendorf, Germany). Sequencing was performed on the Illumina NovaSeq X Plus sequencing platform with paired-end 150 bp cycles at the Institute for Genomic Medicine (IGM), UC San Diego with a target sequencing depth of approximately 10 million read pairs per samples.

### Quality control, taxonomic profiling, antimicrobial resistance gene identification and functional potential profiling

The open-source CZ ID pipeline (https://czid.org/) was used for quality control processing, the taxonomic classification of microbes (mNGS pipeline version 8.3), and antimicrobial resistance genes (AMR pipeline version 1.4.2; CARD Database version 3.2.6; Wildcard Database version 4.0.0). CZ ID pipeline data pre-processing involved subtractive alignment to mouse host genome (Bowtie2 and Hisat2) using raw .fastq files and fastq quality control^47^. Prior to quality control, processing samples had an average number of 21,831,292 ± 9,115,324 reads. After quality control, samples had an average number of 12,192,985 ± 6,484,534 reads (**Supplemental Table 2**).

For taxonomic classification, microbial reads were aligned to the NCBI NT database (National Center for Biotechnology Information Nucleotide Database) using minimap2 (assembly-based alignment). Microbial reads were also aligned to the NCBI NR database (non-redundant protein) using Diamond. Taxa were retained for downstream analysis if they met all of the following criteria: an average NT alignment length (L) > 50 bp, an NT abundance > 10 reads per mission (rPM), and at least one aligning read in both the NT and NR databases (NT r > 0 and NR r > 0) to reduce spurious hits. To represent each sample as a proportional composition of its detected taxa, we converted NT rPM table from CZ ID into relative abundances by dividing each taxon’s NT rPM by the sample’s total NT rPM.

In parallel with taxonomic profiling, samples were run through CZ ID’s AMR pipeline (Comprehensive Antibiotic Resistance Database Resistance Gene Identifier tool) and antimicrobial resistance genes that and ≥ 5% read coverage breadth were kept for resistance analyses.

Functional pathway potential profiling was conducted using the bioBakery WGS pipeline (UniRef90; KEGG Orthology (KO)) implemented in Nephele^48^. Paired-end .fastq files generated by CZ ID pipeline following its mouse host filtering step were used as input, since the Nephele bioBakery pipeline does not include a mouse reference for host-read removal.

### Statistical analysis

All downstream analyses were performed in R (version 4.6.0) using custom scripts. Taxonomic and ARG feature tables were constructed from the CZ ID output and filtered in R. For taxonomic analyses, species-and genus-level bacterial abundances were expressed as relative abundances (NT rPM normalized to the per-sample total). For ARG analyses, gene-level abundances (depth per million, dpm) were similarly converted to relative abundances. Functional potential profiling used HUMAnN3 pathway abundances (copies per million, cpm) converted to relative abundance. All diversity and differential abundance analyses were stratified (performed separately within each host sex x treatment group).

Alpha diversity was assessed using Shannon diversity index, calculated with vegan package (version 2.8-0) on the relative-abundance feature table. Differences in Shannon diversity between stool and culture samples were evaluated within each sex x treatment stratum using paired t-tests. For ARGs, total ARG burden (summed dpm per sample) and ARG richness (number of distinct genes with non-zero abundance per sample) were compared between sample types using the same paired approach.

Beta diversity was assessed using Bray-Curtis dissimilarities computed with vegan on the relative-abundance feature table. Differences in community composition between sample types were tested using permutational multivariate analysis of variance (PERMANOVA; adonis2, vegan package), with permutations accounting for repeated sampling of the same mouse. Homogeneity of multivariate dispersion was assessed using PERMDISP (betadisper, vegan package) to confirm that PERMANOVA results were not attributable to differences in within-group dispersion. Community structure was visualized by principal coordinate analysis (PCoA) of the Bray-Curtis dissimilarity matrix using classical multidimensional scaling (cmdscale, stats package), with the first two coordinates retained for plotting.

Differentially abundant taxa, ARGs, and functional potential pathways were identified using MaAsLin2 (version 1.26.0)^23^. Feature tables were already expressed as relative abundances so no additional normalization was applied and features were log-transformed. Models included sample type as the fixed effect of interest and replicate ID as a random effect to account for repeated measures from the same mouse; a minimum prevalence threshold of 0.05 was applied. Statistical significance was assessed at a Benjamini-Hochberg false discovery rate-corrected q-value < 0.05. For sex comparisons, host sex was specified as the fixed effect of interest.

Differential abundance analysis results were visualized as volcano plots of the MaAsLin2 coefficient against the -log10 q-value, and as heatmaps of significant features using ComplexHeatmap (version 2.28.0); for heatmaps, relative abundances were Z-score-scaled across samples. Taxonomic composition was additionally summarized using stacked relative-abundance bar plots. Figures were generated using ggplot2 (version 4.0.3).

## Acknowledgements

This work was supported by resources at the Center for Microbiome Innovation, the UC San Diego animal care facilities, and the professional veterinary staff for providing their valuable services. This work includes data generated at the UC San Diego IGM Genomics Center utilizing an Illumina NovaSeq X Plus that was purchased with funding from a National Institutes of Health SIG grant (#S10 OD026929).

## Contributions

**Weimiao Long:** Conceptualization; investigation; writing – original draft; writing – review and editing. **Avani Tantry**: Investigation. **Julielam Tran**: Investigation. **Erika L. Cyphert**: Conceptualization; validation; investigation; writing – original draft; writing – review and editing; supervision; project administration.

## Data Availability

Raw shotgun metagenomic sequencing .fastq files are available at Sequence Read Archive accession ID PRJNA1504325.

## References

1. E Thursby & N Juge. Introduction to the human gut microbiota. Biochem J 474, 1823– 1836 (2017).

2. FC Pereira & D Berry. Microbial nutrient niches in the gut. Env. Microbiol 19, 1366– 1378 (2017).

3. C Nowicki et al. Comparison of gut microbiome composition in colonic biopsies, endoscopically-collected and at-home-collected stool samples. Front Microbiol 14, (2023).

4. Q Tang et al. Current sampling methods for gut microbiota: a call for more precise devices. Front Cell Infect Microbiol 10, (2020).

5. GR Gibson, JH Cummings, & GT Macfarlane. Use of a three-stage continuous culture system to study the effect of mucin on dissimilatory sulfate reduction and methanogenesis by mixed populations of human gut bacteria. Appl Env. Microbiol 54, 2750–2755 (1988).

6. C Cinquin, G Le Blay, I Fliss, & C Lacroix. New three-stage in vitro model for infant colonic fermentation with immobilized fecal microbiota. FEMS Microbiol Ecol 57, 324–336 (2006).

7. L Liu et al. Establishing a mucosal gut microbial community in vitro using an artificial simulator. PLoS One 13, e0197692 (2018).

8. Z Jin, A Ng, CF Maurice, & D Juncker. The mini colon model: a benchtop multi-bioreactor system to investigate the gut microbiome. Gut Microbes 14, 2096993 (2022).

9. S Ahmadi et al. An in vitro batch-culture model to estimate the efects of interventional regimens on human fecal microbiota. J Vis Exp 149, e59524 (2019).

10. A Aranda-Diaz et al. Establishment and characterization of stable, diverse, fecal-derived in vitro microbial communities that model the intestinal microbiota. Cell Host Microbe 30, 260– 272 (2022).

11. L Yang et al. An in vitro evaluation of the effect of Bifidobacterium longum L556 on microbiota composition and metabolic properties in patients with coronary heart disease (CHD). Probiotics Antimicrob Proteins 17, 3184–3198 (2025).

12. F Yousi et al. Evaluation of the effects of four media on human intestinal microbiota culture in vitro. AMB Express 9, 69 (2019).

13. P Shah et al. A microfluidics-based in vitro model of the gastrointestinal human-microbe interface. Nat Commun 7, 11535 (2016).

14. M Calvigioni et al. Development of an in vitro model of the gut microbiota enriched in mucus-adhering bacteria. Microbiol Spectr 11, e00336–23 (2023).

15. HJ Kim, H Li, JJ Collins, & DE Ingber. Contributions of microbiome and mechanical deformation to intestinal bacterial overgrowth and inflammation in a human gut-on-a-chip. PNAS 113, E7–15 (2016).

16. DA Goldman, et al. Competition for shared resources increases dependence on initial population size during coalescence of gut microbial communities. PNAS 122, e2322440122 (2025).

17. AI Celis, DA Relman, & KC Huang. The impact of iron and heme availability on the healthy human gut microbiome in vivo and in vitro. Cell Chem Biol 30, 110–126 (2023).

18. RK Dudek-Wicher, A Junka, & M Bartoszewicz. The influence of antibiotics and dietary components on gut microbiota. Prz Gastroenterol 13, 85–92 (2018).

19. J Zhang, L Chen, A Gomez-Simmonds, MT Yin, & DE Freedberg. Antibiotic-specific risk for community-acquired Clostridioides difficile infection in the United States from 2008 to 2020. Antimicrob Agents Chemother 66, e01129–22 (2022).

20. DV Patangia, CA Ryan, E Dempsey, RP Ross, & C Stanton. Impact of antibiotics on the human microbiome and consequences for host health. Microbiologyopen 11, e1260 (2022).

21. Z Chaudhry, A Cazares, N Thomson, & RS Heyderman. Profiling the gut resistome to unlock antimicrobial resistance biology and inform clinical risk. Nat Comm 17, 6212 (2026).

22. KF Sutton & LW Ashley. Antimicrobial resistance in the United States: Origins and future directions. Epidemiol Infect 152, e33 (2024).

23. H Mallick et al. Multivariable association discovery in population-scale meta-omics studies. PLoS Comput Biol 17, e1009442 (2021).

24. F Beghini et al. Integrating taxonomic, functional, and strain-level profiling of diverse microbial communities with bioBakery 3. eLife 10, e65088 (2021).

25. MA Schell et al. The genome sequence of Bifidobacterium longum reflects its adaptation to the human gastrointestinal tract. PNAS 99, 14422–14427 (2002).

26. BM Woodall et al. Enterococcus faecalis readily adapts membrane phospholipid composition to environmental and genetic perturbation. Front Microbiol 12, (2021).

27. F Depardieu et al. New combinations of mutations in vanD-type vancomycin-resistant Enterococcus faecium, Enterococcus faecalis, and Enterococcus avium strains. Antimicrob Agents Chemother 53, 1952–1963 (2009).

28. HB Kim et al. oqxAB encoding a multidrug efflux pump in human clinical isolates of Enterobacteriaceae. Antimicrob Agents Chemother 53, 3582–3584 (2009).

29. EL Cyphert et al. Commensal taxa in gut microbiota limit antibiotic resistance during extended oral antibiotic use. bioRxiv 10.1101/2025.08.13.670183 (2025) doi:10.1101/2025.08.13.670183.

30. H Juricova, J Matiasovicova, T Kubasova, D Cejkova, & I Rychlik. The distribution of antibiotic resistance genes in chicken gut microbiota commensals. Sci Rep 11, 3290 (2021).

31. TM Coque, KV Singh, GM Weinstock, & BE Murray. Characterization of dihydrofolate reductase genes from trimethoprim-susceptible and trimethoprim-resistant strains of Enterococcus faecalis. Antimicrob Agents Chemother 43, 141–147 (1999).

32. EW Lee, MN Huda, T Kuroda, T Mizushima, & T Tsuchiya. EfrAB, an ABC multidrug efflux pump in Enterococcus faecalis. Antimicrob Agents Chemother 47, 3733–3738 (2003).

33. KV Singh, GM Weinstock, & BE Murray. An Enterococcus faecalis ABC homologue (Lsa) is required for the resistance of this species to clindamycin and quinupristin-dalfopristin. Antimicrob Agents Chemother 46, 1845–1850 (2002).

34. EW Lee et al. Functional cloning and expression of emeA, and characterization of emeA, a multidrug efflux pump from Enterococcus faecalis. Biol Pharm Bull 26, 266–270 (2003).

35. F Serafini et al. Insights into physiological and genetic mupirocin susceptibility in Bifidobacteria. Appl Env. Microbiol 77, 3141–3146 (2011).

36. BJ Kim, HY Kim, YJ Yun, BJ Kim, & YH Kook. Differentiation of Bifidobacterium species using partial RNA polymerase beta-subunit (rpoB) gene sequences. Int J Syst Evol Microbiol 60, 2697–2704 (2010).

37. EJ Culp & AL Goodman. Cross-feeding in the gut microbiome: ecology and mechanisms. Cell Host Microbe 31, 485–499 (2023).

38. Y Zhu et al. Exploration of the Muribaculaceae family in the gut microbiota: diversity, metabolism, and function. Nutrients 16, 2660 (2024).

39. RS Gupta, A Nanda, & B Khadka. Novel molecular, structural and evolutionary characteristics of the phosphoketolases from Bifidobacteria and Coriobacteriales. PLoS One 12, e0172176 (2017).

40. M Centanni et al. Bifidobacterium pseudolongum in the ceca of rats fed hi-maize starch has characteristics of a keystone species in Bifidobacterial blooms. Appl Env. Microbiol 84, e00547–18 (2018).

41. M Derrien, EE Vaughan, CM Plugge, & WM de Vos. Akkermansia muciniphila gen. nov., sp. nov., a human intestinal mucin-degrading bacterium. Int J Sys Evol Microbiol 54, 1469–1476 (2004).

42. A Aranda-Diaz et al. Assembly of stool-derived bacterial communities follows ‘early-bird’ resource utilization dynamics. Cell Syst 16, 101240 (2025).

43. A Zarrinpar et al. Antibiotic-induced microbiome depletion alters metabolic homeostasis by affecting gut signaling and colonic metabolism. Nat Comm 9, 2872 (2018).

44. C Allaband et al. Time of sample collection is critical for the replicability of microbiome analyses. Nat Metab 6, 1282–1293 (2024).

45. R Sender, S Fuchs, & R Milo. Revised estimates for the number of human and bacteria cells in the body. PLoS Biol 14, e1002533 (2016).

46. C Marotz et al. SARS-CoV-2 detection status associates with bacterial community composition in patients and the hospital environment. Microbiome 9, 132 (2021).

47. KL Kalantar et al. IDseq - An open source cloud-based pipeline and analysis service for metagenomic pathogen detection and monitoring. GigaScience 9, giaa111 (2020).

48. N Weber et al. Nephele: a cloud platform for simplified, standardized and reproducible microbiome data analysis. Bioinformatics 34, 1411–1413 (2018).

