## Supplementary Figures for "Validating a high-throughput *in vitro* model for culturing antibiotic-altered microbiome communities"

**Running Title:** Validating model for culturing altered microbiota

**Supplemental Table 1.** Mapping function from Metacyc class to a high-level ontology.

| **MetaCyc class** | **High-level ontology** |
| --- | --- |
| Energy-Metabolism | Energy metabolism |
| C1-COMPOUNDS |  |
| Cofactor-Biosynthesis | Cofactor biosynthesis |
| Tetrapyrrole-Biosynthesis |  |
| Polyprenyl-Biosynthesis |  |
| Amino-Acid-Biosynthesis | Amino acid metabolism |
| Amino-Acid-Degradation |  |
| Aminoacyl-tRNAs-Charging |  |
| AMINE-DEG |  |
| Polyamine-Biosynthesis |  |
| Carbohydrates-Degradation | Carbohydrate metabolism |
| Carbohydrates-Biosynthesis |  |
| Glycan-Pathways |  |
| Polymer-Degradation |  |
| CYCLITOLS-DEG |  |
| Cell-Structure-Biosynthesis |  |
| Nucleotide-Biosynthesis | Nucleotide metabolism |
| NUCLEO-DEG |  |
| Nucleic-Acid-Processing |  |
| Lipid-Biosynthesis | Lipid metabolism |
| Fatty-Acid-and-Lipid-Degradation |  |
| *anything else* | Other |

**Supplemental Table 2.** Quality control processing of shotgun metagenomic paired-end samples using the Chan Zuckerberg ID pipeline.

| Sample name | Total reads (combined paired-end) | Reads after bowtie2 ercc filtered | Reads after fastp | Reads after bowtie2 host filtered | Reads after hisat2 host filtered | Reads after czid dedup |
| --- | --- | --- | --- | --- | --- | --- |
| 25_S25 | 23410056 | 23410006 | 21878790 | 21548850 | 21548812 | 19692350 |
| 26_S26 | 12255852 | 12255850 | 11137866 | 11133024 | 11133024 | 10639820 |
| 27_S27 | 18239928 | 18239924 | 16386028 | 16372568 | 16372566 | 15176802 |
| 28_S28 | 35233326 | 35233298 | 31606992 | 31418594 | 31418530 | 26695158 |
| 29_S29 | 21694886 | 21694836 | 18510988 | 17886624 | 17886520 | 15669922 |
| 30_S30 | 16548714 | 16548674 | 14832548 | 14358170 | 14358134 | 13009470 |
| 31_S31 | 13619204 | 13619190 | 12882156 | 12584840 | 12584822 | 11567282 |
| 32_S32 | 18371646 | 18371632 | 17356790 | 16921372 | 16921334 | 15532986 |
| 33_S33 | 15849850 | 15849850 | 14904978 | 14897268 | 14897268 | 14219744 |
| 34_S34 | 20887074 | 20887064 | 19193216 | 19162884 | 19162880 | 17436196 |
| 35_S35 | 15179852 | 15179848 | 14317118 | 14308140 | 14308140 | 13437148 |
| 36_S36 | 16725646 | 16725642 | 15285822 | 15257774 | 15257764 | 13777200 |
| 37_S37 | 19755244 | 19755186 | 18439446 | 17380380 | 17380308 | 15815516 |
| 38_S38 | 14402076 | 14402058 | 13326886 | 13025204 | 13025190 | 12084056 |
| 39_S39 | 16939014 | 16938996 | 15843418 | 15517956 | 15517932 | 14256134 |
| 40_S49 | 27749428 | 27749426 | 26058070 | 26057018 | 26057018 | 23357668 |
| 41_S50 | 17134092 | 17134090 | 16516664 | 16516086 | 16516086 | 15069158 |
| 42_S51 | 23465672 | 23465670 | 22157772 | 22156526 | 22156526 | 19855688 |
| 43_S52 | 17088212 | 17088204 | 14663222 | 14662540 | 14662540 | 13899756 |
| 44_S53 | 21718904 | 21718904 | 20939576 | 20938216 | 20938216 | 9014904 |
| 45_S54 | 21979134 | 21979134 | 21024844 | 21022110 | 21022110 | 4289256 |
| 46_S55 | 38635528 | 38635528 | 33717836 | 33714646 | 33714646 | 7997236 |
| 47_S56 | 29000116 | 29000114 | 26670278 | 26668544 | 26668544 | 10387030 |
| 48_S57 | 13349486 | 13349486 | 12864164 | 12863728 | 12863728 | 11578270 |
| 49_S58 | 12765332 | 12765332 | 9578068 | 9577536 | 9577536 | 9139798 |
| 50_S59 | 30467630 | 30467630 | 29104022 | 29102136 | 29102136 | 25480306 |
| 51_S60 | 25758118 | 25758118 | 24366982 | 24364142 | 24364142 | 7477796 |
| 52_S61 | 18248972 | 18248970 | 17526068 | 17523172 | 17523172 | 4628216 |
| 53_S62 | 27951412 | 27951410 | 26037632 | 26035764 | 26035764 | 6129910 |
| 54_S63 | 18609990 | 18609990 | 16715962 | 16713794 | 16713794 | 2166898 |
| 55_S64 | 14432414 | 14432414 | 13880218 | 13879478 | 13879478 | 11446314 |
| 56_S65 | 27045696 | 27045692 | 26031066 | 26029950 | 26029950 | 22843508 |
| 57_S66 | 19920640 | 19920614 | 18823218 | 18822092 | 18822092 | 17035806 |
| 58_S67 | 18223002 | 18222998 | 17225394 | 17224448 | 17224448 | 15431730 |
| 59_S68 | 24794844 | 24794836 | 23676584 | 23675562 | 23675562 | 10023860 |
| 60_S69 | 23796078 | 23796078 | 22168196 | 22165942 | 22165942 | 7569282 |
| 61_S70 | 55222268 | 55222264 | 48535842 | 48531444 | 48531444 | 7619430 |
| 62_S71 | 12807872 | 12807872 | 12240458 | 12239678 | 12239678 | 4714706 |
| 63_S72 | 25489192 | 25489192 | 24513764 | 24512318 | 24512318 | 21612750 |
| 64_S73 | 16934038 | 16933982 | 16319900 | 16318940 | 16318940 | 5766750 |
| 65_S74 | 22775842 | 22775842 | 21672686 | 21671142 | 21671142 | 18986498 |
| 66_S75 | 18931040 | 18931040 | 18156434 | 18154136 | 18154136 | 8005452 |
| 67_S76 | 29963874 | 29963872 | 27919560 | 27916090 | 27916090 | 9880252 |
| 68_S77 | 60877504 | 60877504 | 48102350 | 48098060 | 48098060 | 9354880 |
| 69_S78 | 21528494 | 21528494 | 20082794 | 20080492 | 20080492 | 4023778 |
| 70_S79 | 21220848 | 21220838 | 20074480 | 20073334 | 20073334 | 17603170 |
| 71_S80 | 25234186 | 25234180 | 23934526 | 23933480 | 23933480 | 20780590 |
| 72_S81 | 21501110 | 21501100 | 20098054 | 20096674 | 20096674 | 18352842 |
| 73_S82 | 18168076 | 18168070 | 17266954 | 17266094 | 17266094 | 15956372 |
| 74_S83 | 17704336 | 17704322 | 16919346 | 16918048 | 16918048 | 6676434 |
| 75_S84 | 16647808 | 16647808 | 15886302 | 15883940 | 15883940 | 4700328 |
| 76_S85 | 16180870 | 16180870 | 14988836 | 14986330 | 14986330 | 2772330 |
| 77_S86 | 27359790 | 27359790 | 26250616 | 26248320 | 26248320 | 8948936 |
| 78_S87 | 2563636 | 2563636 | 538566 | 538542 | 538542 | 513730 |
| 79_S88 | 20718018 | 20717984 | 19479280 | 19478012 | 19478012 | 16638180 |
| 80_S89 | 32131016 | 32131014 | 30452426 | 30450304 | 30450304 | 25866234 |
| 81_S90 | 19180492 | 19180488 | 18452428 | 18448204 | 18448204 | 4153676 |
| 82_S91 | 18686002 | 18686002 | 17959022 | 17955568 | 17955568 | 5902232 |
| 83_S92 | 17716522 | 17716522 | 16904488 | 16902656 | 16902656 | 5471372 |
| 84_S93 | 19087640 | 19087640 | 17970212 | 17967390 | 17967390 | 3446012 |

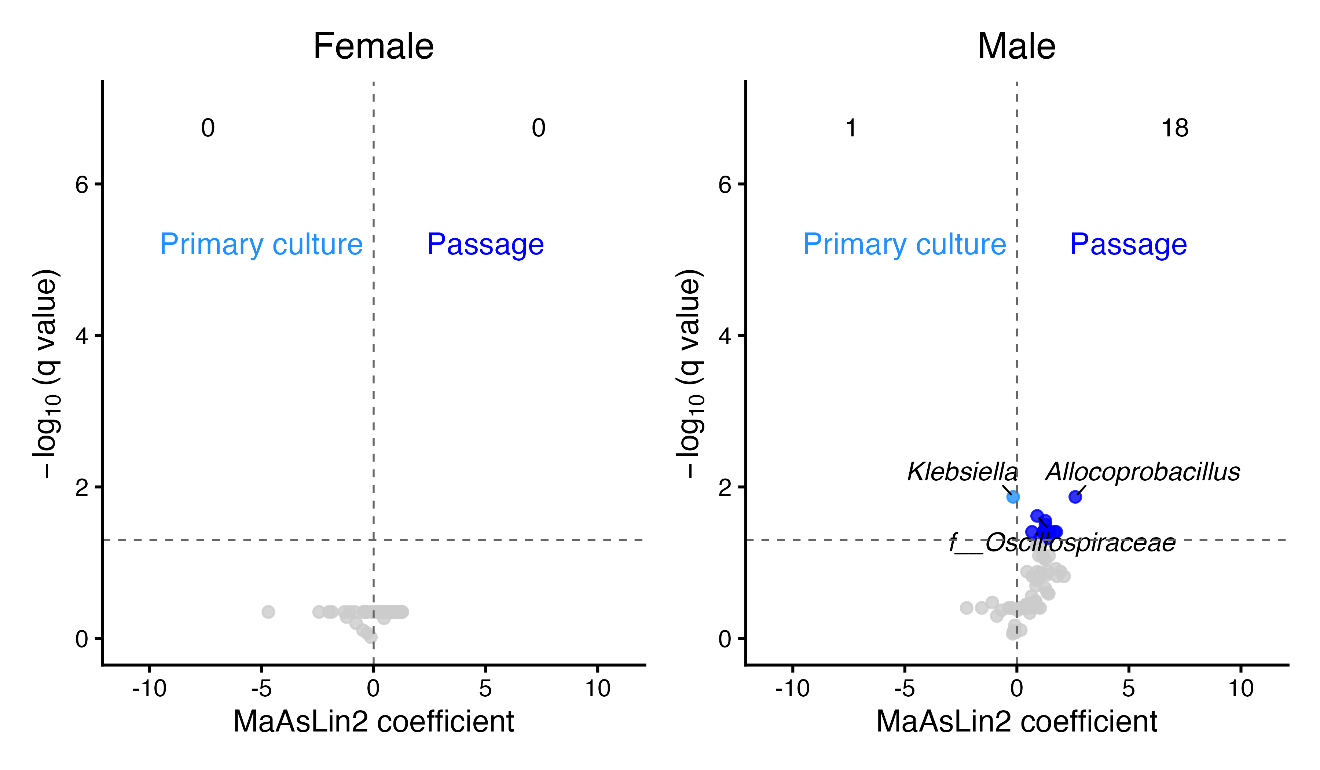

**Supplemental Figure 1**. Volcano plots of differentially abundant genera between primary culture and passage in the Female (left) and Male (right) Altered groups. Genera significantly enriched in primary culture (q < 0.05) are in light blue, those significantly enriched in culture are in dark blue, and non-significant genera are in grey.

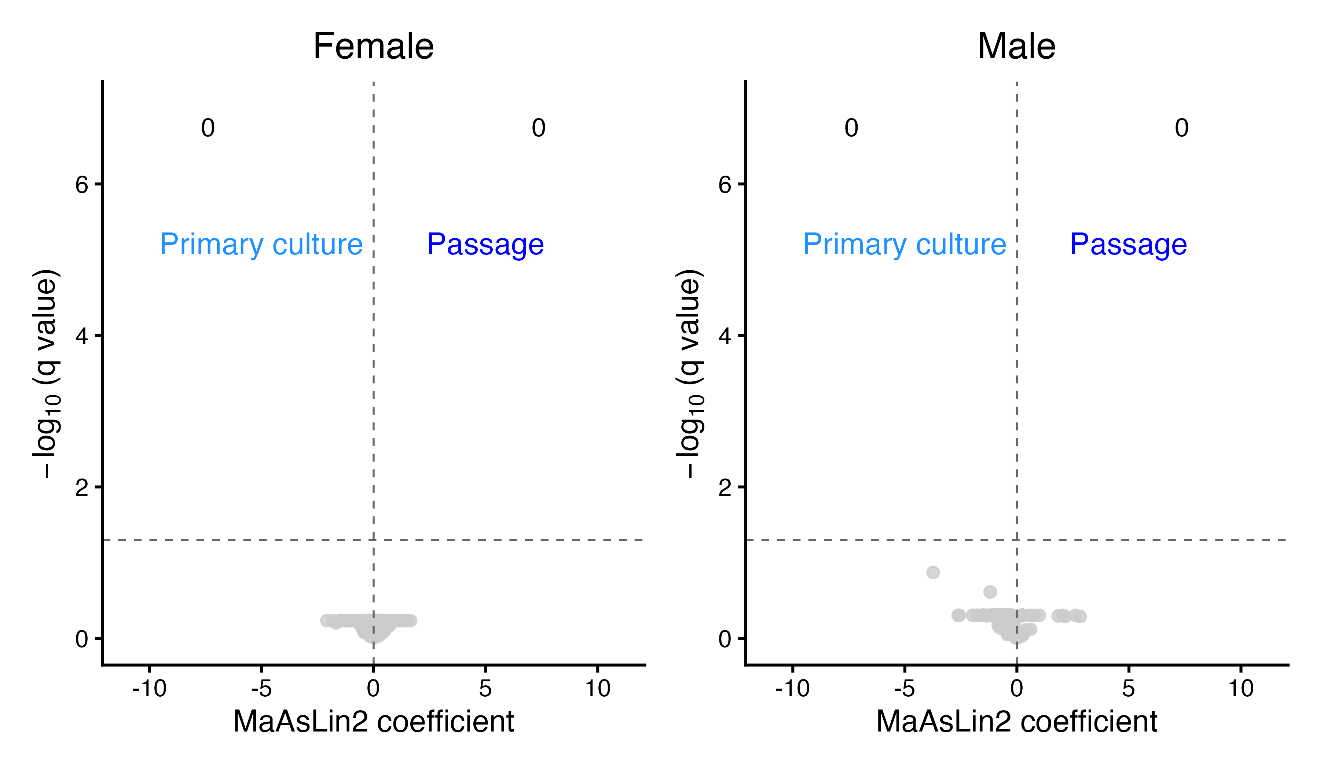

**Supplemental Figure 2.** Volcano plots of differentially abundant genera between primary culture and passage in the Female (left) and Male (right) Unaltered groups. Genera significantly enriched in primary culture (q < 0.05) are in light blue, those significantly enriched in culture are in dark blue, and non-significant genera are in grey.

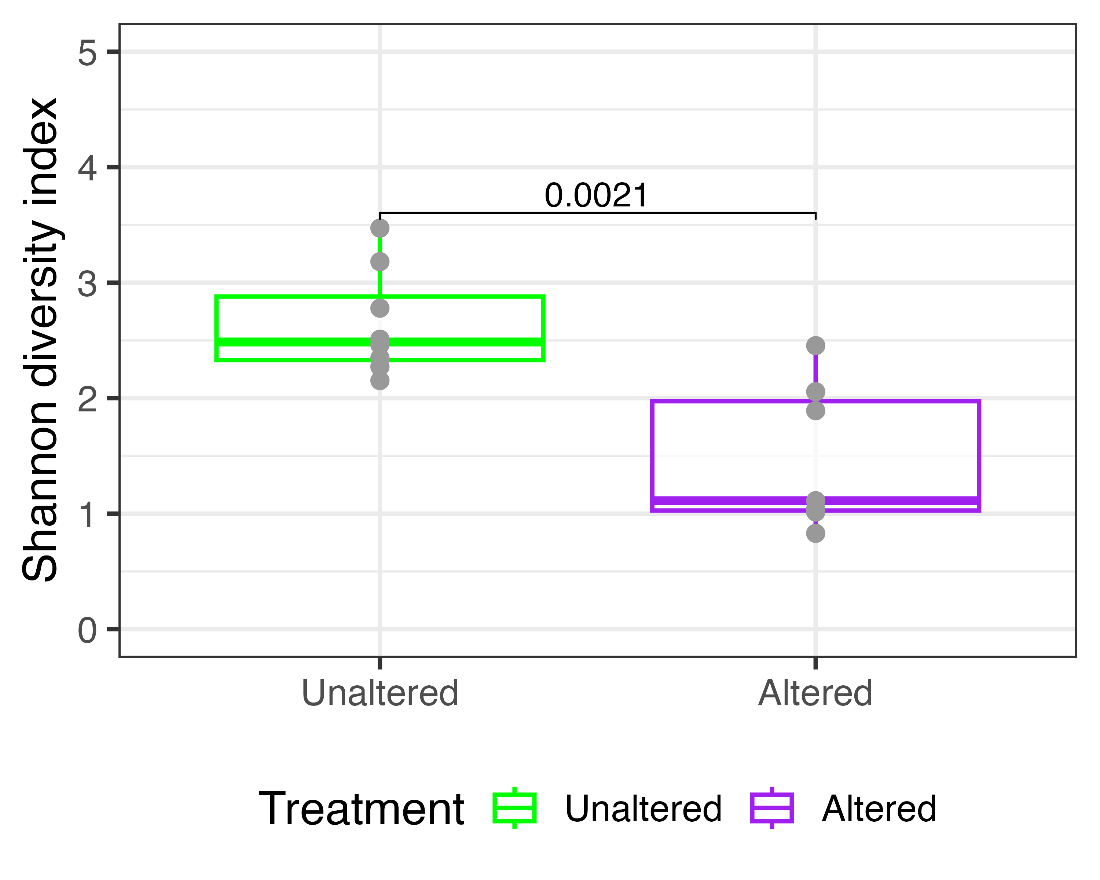

**Supplemental Figure 3.** Boxplot comparing Shannon diversity index of genus level composition between Unaltered stool microbiome and Altered stool microbiome pooling both sexes. A Welch (two-sample) T-test was used to evaluate statistical significance.

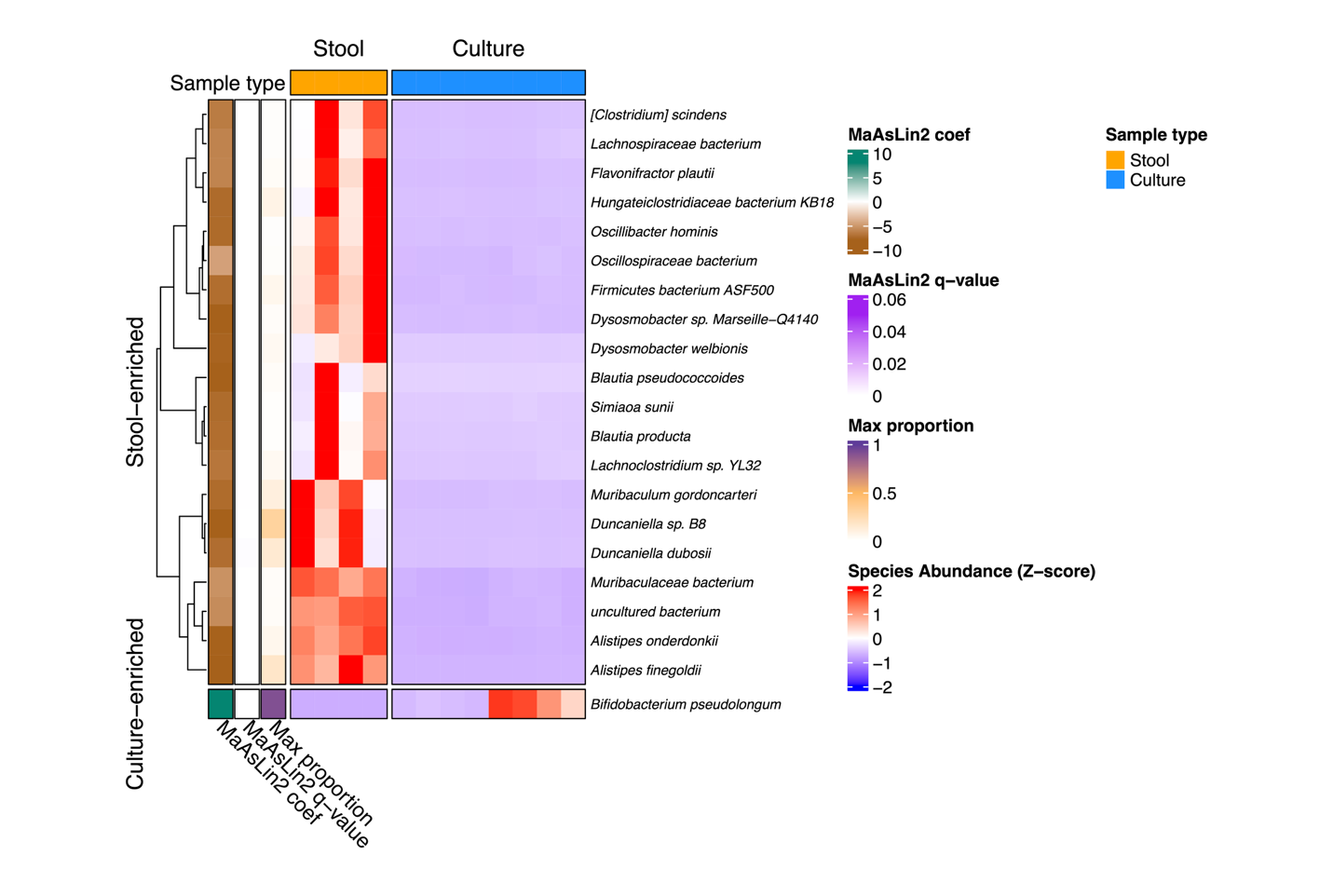

**Supplemental Figure 4.** Heatmap of differentially enriched species between stool and culture in Female Unaltered group. Feature values were normalized using row-wise Z score transformation (mean = 0, standard deviation = 1) to emphasize relative differences across samples. Rows represent the genera that is significantly enriched in either group (MaAsLin2 q-value < 0.05; Max proportion > 1%); columns represent individual samples with annotated metadata shown above.

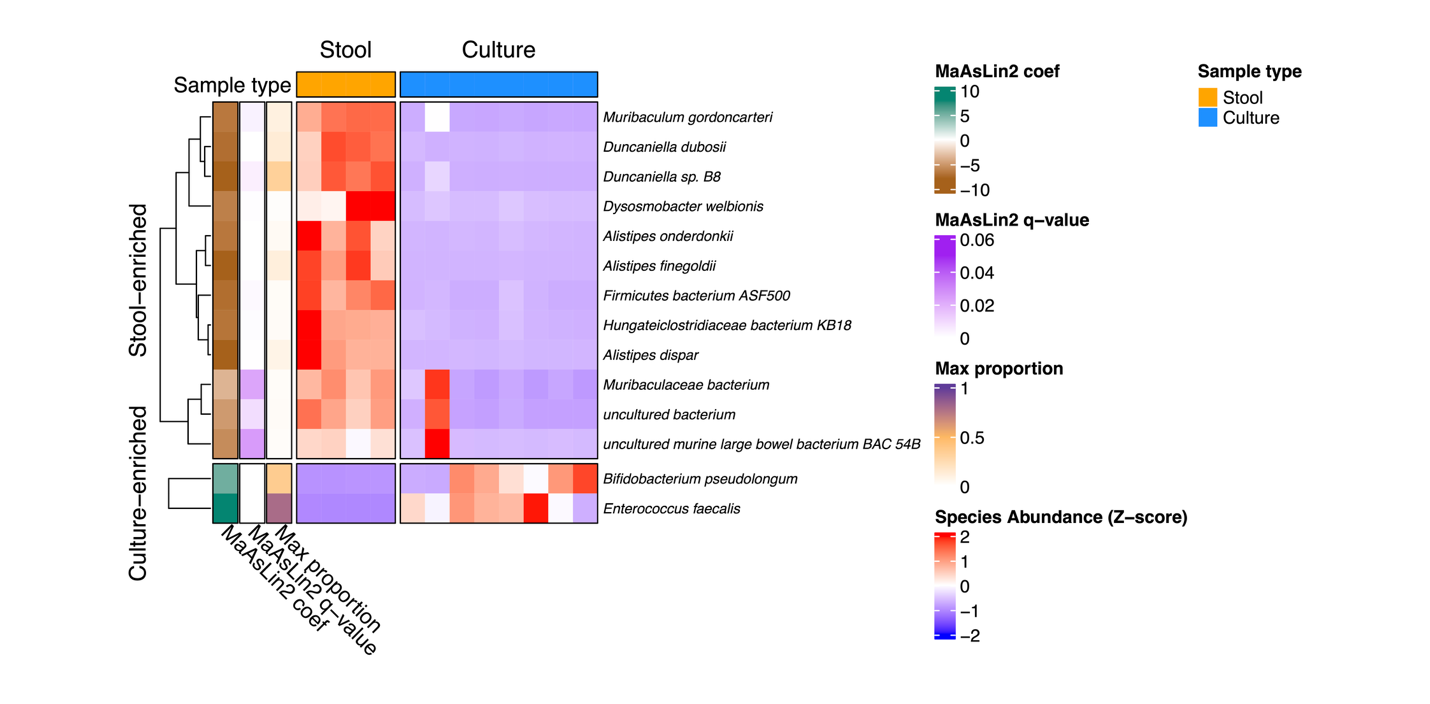

**Supplemental Figure 5.** Heatmap of differentially enriched species between stool and culture in Male Unaltered group. Heatmap construction and annotations are as described above.
